# The lectin LecB induces patches with basolateral characteristics at the apical membrane to promote *Pseudomonas aeruginosa* host cell invasion

**DOI:** 10.1101/2022.03.23.485577

**Authors:** Roland Thuenauer, Katja Kühn, Yubing Guo, Fruzsina Kotsis, Maokai Xu, Anne Trefzer, Silke Altmann, Sarah Wehrum, Najmeh Heshmatpour, Brian Faust, Alessia Landi, Britta Diedrich, Jörn Dengjel, E. Wolfgang Kuehn, Anne Imberty, Winfried Römer

## Abstract

The opportunistic bacterium *Pseudomonas aeruginosa* can infect mucosal tissues of the human body. To persist at the mucosal barrier, this highly adaptable pathogen has evolved many strategies, including invasion of host cells. Here, we show that the *P. aeruginosa* lectin LecB binds and cross-links fucosylated receptors at the apical plasma membrane of epithelial cells. This triggers a signaling cascade via Src kinases and PI3K leading to the formation of patches enriched with the basolateral marker PIP_3_ at the apical plasma membrane. This identifies LecB as causative bacterial factor for activating this well-known host cell response that is elicited upon apical binding of *P. aeruginosa*. Downstream of PI3K, Rac1 is activated to cause actin rearrangement and the outgrowth of protrusions at the apical plasma membrane. LecB-triggered PI3K activation also results in aberrant recruitment of caveolin-1 to the apical domain. In addition, we reveal a positive feedback loop between PI3K activation and apical caveolin-1 recruitment, which provides a mechanistic explanation for the previously observed implication of caveolin-1 in *P. aeruginosa* host cell invasion. Interestingly, LecB treatment also reversibly removes primary cilia. To directly prove the role of LecB for bacterial uptake, we coated bacteria-sized beads with LecB, which drastically enhanced their endocytosis. Furthermore, LecB deletion and LecB inhibition with L-fucose diminished the invasion efficiency of *P. aeruginosa* bacteria. Taken together, our study identifies LecB as missing link that can explain how PI3K signaling and caveolin-1 recruitment are triggered to facilitate invasion of epithelial cells from the apical side by *P. aeruginosa*.

**Importance:** An intriguing feature of the bacterium *P. aeruginosa* is its ability to colonize highly diverse niches. *P. aeruginosa* can, besides biofilm formation, also enter and proliferate within epithelial host cells. Moreover, research during recent years has shown that *P. aeruginosa* possesses many different mechanisms to invade host cells. In this study we identify LecB as novel invasion factor. In particular, we show that LecB activates PI3K signaling, which is connected via a positive feedback loop to apical caveolin-1 recruitment, and leads to actin rearrangement at the apical plasma membrane. This provides a unifying explanation for the previously reported implication of PI3K and caveolin-1 in *P. aeruginosa* host cell invasion. In addition, our study adds a further function to the remarkable repertoire of the lectin LecB, which is all brought about by the capability of LecB to recognize fucosylated glycans on many different niche-specific host cell receptors.

## Introduction

*Pseudomonas aeruginosa* is a ubiquitous environmental bacterium. Due to its intrinsic adaptability and the rise of multi-drug resistant strains this bacterium poses a dangerous threat especially in hospital settings. Accordingly, carbapenem-resistant *P. aeruginosa* strains were categorized by the World Health Organization (WHO) as priority 1 pathogens for which new antibiotics are critically required (1).

When infecting a human host, *P. aeruginosa* can switch between many lifestyles including planktonic behavior and biofilm formation. In addition, evidence accumulated during recent years that *P. aeruginosa* can also invade host cells. It has been demonstrated that *P. aeruginosa* is able to enter and survive (2, 3), move (4), and proliferate (5) in non-phagocytic cells. Moreover, after being taken up by macrophages, *P. aeruginosa* can escape phagosomes, and eventually lyse macrophages from the inside (6). The importance of the intracellular lifestyle for *P. aeruginosa* is supported by the observation that this bacterium has a whole arsenal of mechanisms to facilitate uptake by host cells. *P. aeruginosa* can invade by binding the remains of dead cells that are then taken up by surrounding cells through efferocytosis (7), by deploying the effector VgrG2b via its type VI secretion system (T6SS) to promote microtubule-dependent uptake (8), by utilizing cystic fibrosis transmembrane conductance regulator (CFTR) to stimulate caveolin-1 – dependent endocytosis (9), and by interaction between the *P. aeruginosa* lectin LecA and the host cell glycosphingolipid globotriaosylceramide (Gb3) to facilitate invasion through a lipid zipper mechanism (10).

After incorporation by a human host, *P. aeruginosa* will typically interact with the apical plasma membranes of epithelial cells lining the mucosae. Interestingly, *P. aeruginosa* has developed mechanisms to manipulate the apical identity of these membranes. The hallmark of this process is the activation of phosphatidylinositol 3-kinase (PI3K) resulting in abnormal accumulation of phosphatidylinositol (3,4,5)-trisphosphate (PIP_3_) at the apical plasma membrane, which eventually generates patches with basolateral characteristics at the apical plasma membrane (11). This inversion of polarity has been suggested to help in binding of *P. aeruginosa* to cells since this bacterium uses different mechanisms to bind apical and basolateral plasma membranes (12). It is also crucial for host cell invasion by *P. aeruginosa*, because inhibition of PI3K signaling markedly reduces bacterial uptake (13). However, the exact mechanism how *P. aeruginosa* is able to convert apical to basolateral plasma membrane is not clear. The formation of patches with basolateral characteristics at the apical plasma membrane requires the type III secretion system (T3SS), but strikingly does not require any of the toxins that are secreted via the T3SS (14, 15). To explain these observations two hypotheses were suggested: Host cell membrane damage through bacteria might be the initial event leading to basolateral patch formation, or PI3K signaling is triggered by a still unknown factor from *P. aeruginosa* (11).

Here, we provide data showing that the tetrameric fucose-specific lectin LecB (16), which is exposed at the outer membrane of *P. aeruginosa* (17, 18), represents the missing link. We showed already in a previous publication that purified LecB is able to bind receptors at the apical and basolateral plasma membrane of polarized Madin-Darby canine kidney (MDCK) cells (19). On the basolateral side LecB was able to bind integrins, which led to integrin internalization and loss of epithelial polarity. Since only minute amounts of integrins are found at the apical side of polarized MDCK cells (19, 20), LecB did not dissolve epithelial polarity when applied only to the apical side (19). Here we reveal that binding of LecB to fucosylated apical receptors on epithelial host cells was sufficient to trigger a different signaling cascade in order to promote cellular uptake of *P. aeruginosa*. Apical LecB binding led to Src signaling followed by local PI3K activation, PIP_3_ patch formation at the apical plasma membrane, Rac1 signaling, and actin rearrangement to trigger the formation of protrusions in order to enable host cell invasion of *P. aeruginosa*. In addition, we show that caveolin-1 is recruited abnormally to apical membranes after LecB stimulation and that PI3K activation requires caveolin-1. These data suggest LecB as unifying factor that facilitates and modulates many of the invasion mechanisms that were reported for *P. aeruginosa*.

## Results

### Apical LecB treatment triggers Src-PI3K/Akt signaling

To closer analyze the effects caused by application of purified LecB to the apical side of polarized MDCK cells, we used MDCK cells stably expressing PH-Akt-GFP, which indicates the localization of the lipid PIP_3_ (21) (Fig. 1A). In unstimulated cells, PH-Akt-GFP localized mainly to the basolateral plasma membrane, as expected from the role of PIP_3_ as a basolateral marker in polarized epithelial cells (21). In cells treated apically with LecB, PIP_3_ accumulated at the apical side and protrusions formed that were positive for PH-Akt-GFP (Fig. 1A, white arrows). This replicates the effects that were previously observed after interaction of whole *P. aeruginosa* bacteria with the apical plasma membrane of MDCK cells (22).

**Fig. 1:**
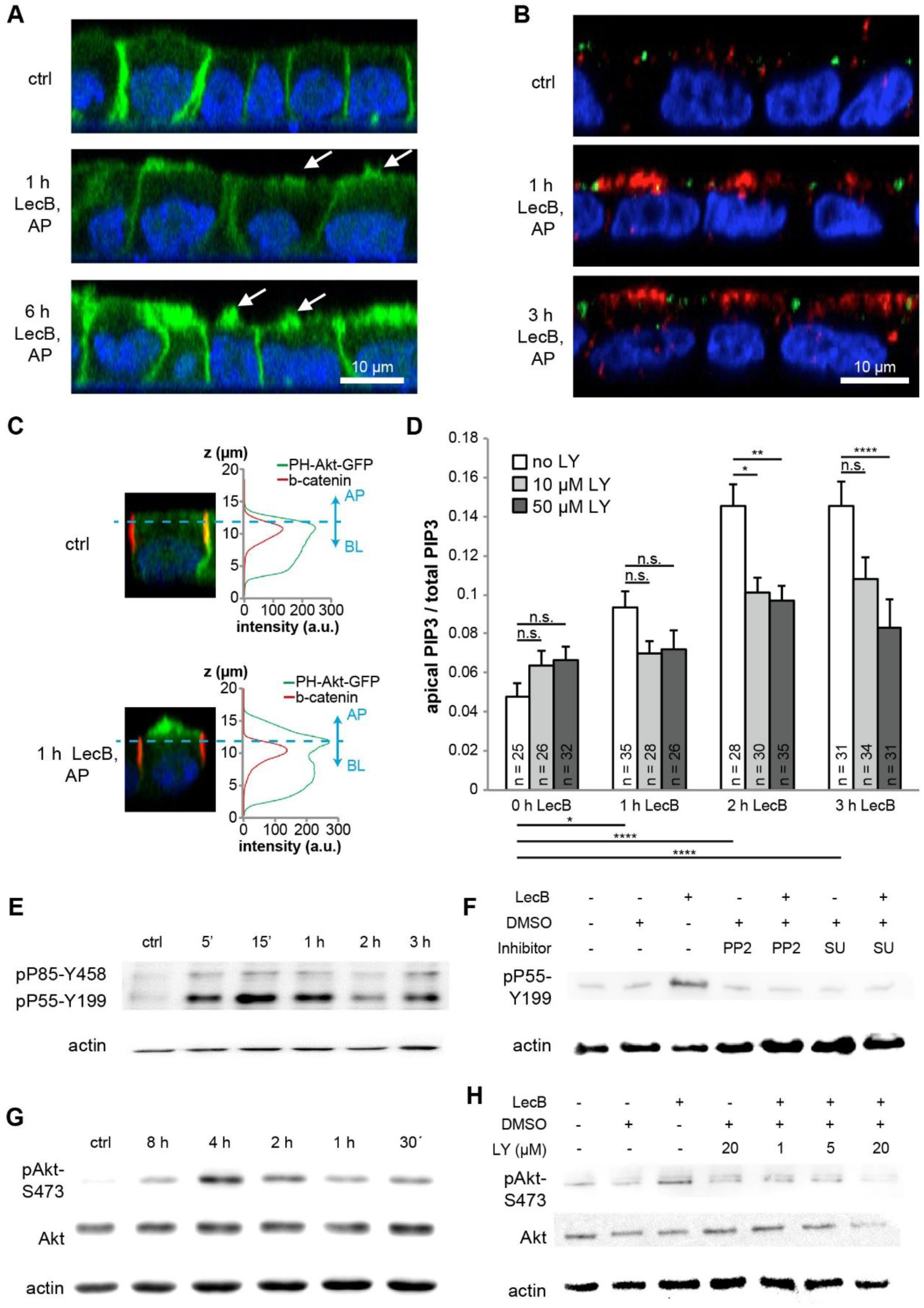
After binding to the apical plasma membrane of MDCK cells, LecB triggers a Src-PI3K/Akt signaling cascade. (A) MDCK cells stably expressing the PIP_3_-marker PH-Akt-GFP (green) were left untreated (ctrl) or treated from the apical (AP) side with LecB for the indicated time periods; nuclei were stained with DAPI (blue). White arrows point to apical protrusion resulting from LecB treatment. (B) MDCK cells were left untreated (ctrl) or treated with LecB as indicated, fixed and then stained for active PI3K (pP85-Y458 and pP55-Y199; red), ZO-1 (green), and nuclei were stained with DAPI (blue). (C) MDCK cells stably expressing the PIP_3_ marker PH-Akt-GFP were treated with LecB from the apical (AP) side, fixed, and stained with β-catenin. To distinguish the apical and basolateral portion of the PH-Akt-GFP signal, β-catenin staining was utilized as ‘ruler’. For the experiment PH-Akt-GFP-positive cells were mixed before seeding with wt cells in a ratio 1:10. This enabled an unbiased quantification by measuring the signals only from PH-Akt-GFP-positive cells that were surrounded by wt cells. (D) Quantification of the experiment described in (C), the numbers indicated at the bottom of each bar represent the number of individual cells that were measured for each condition. Whereas cells treated with LecB show a time-dependent increase of the apical-to-total PH-Akt-GFP/PIP_3_ signal ratio, treatment with LY294002 (LY) reversed this effect. (E) MDCK cells were treated apically with LecB for the indicated times and subjected to WB analysis using an antibody recognizing active PI3K (pP85-Y458 and pP55-Y199). (F) MDCK cells were treated apically with LecB and PP2 (10 µM) or SU6656 (10 µM) for 1 h and subjected to WB analysis utilizing an antibody recognizing active PI3K (pP55-Y199). (G) MDCK cells were treated apically with LecB for the indicated times and subjected to WB analysis utilizing an antibody recognizing active Akt (pAkt-S473). (H) MDCK cells were treated apically with LecB and indicated concentrations of LY294002 (LY) for 1 h and subjected to WB analysis utilizing an antibody recognizing active Akt (pAkt-S473).

Importantly, we demonstrated already previously that apical LecB application did not disturb the integrity of tight junctions (19). Thus, LecB-mediated apical PIP_3_ accumulation cannot be explained by a loss of the barrier function of tight junctions. We therefore investigated if activation of PI3K is the cause of apical PIP3 accumulation. Staining cells with antibodies recognizing active PI3K (pP85-Y458 and pP55-Y199) (23, 24) revealed a clearly visible recruitment and activation of PI3K to subapical regions in LecB-treated cells (Fig. 1B). In addition, incubating the cells with the broad-spectrum PI3K inhibitor LY294002 blocked apical appearance of PH-Akt-GFP after LecB treatment (Figs. 1C and 1D). Activation of PI3K was also detectable by Western Blot (WB) and peaked at approximately 15 min after initiation of LecB stimulation (Fig. 1E). Upstream of PI3K, the activation of Src kinases was required as demonstrated by the ability of the Src kinase inhibitors PP2 and SU6656 to block of LecB-induced PI3K activation (Fig. 1F). LecB activated also Akt, for which phosphorylation at S473 was detectable after 30 min of LecB application and peaked at approximately 4 h (Fig. 1G). Akt signaling occurred downstream of PI3K, because the broad-spectrum PI3K inhibitor LY294002 blocked Akt activation (Fig. 1H). Further tests revealed that the PI3K subunit p110α is mainly responsible for LecB-mediated Akt activation, since the p110α-specific inhibitor PIK-75 blocked Akt activation (Fig. S1A), whereas the p110β-specific inhibitor TGX-221 did not (Fig. S1B). Another fucose-binding lectin, *Ulex europaeus* agglutinin I (UEA-I), failed to replicate LecB-triggered Akt signaling (Fig. S1C), thus indicating that the observed effects are specific for LecB.

To demonstrate that LecB-mediated PI3K/Akt activation is not limited to MDCK cells, we carried out experiments in other cell lines. We chose H1975 lung epithelial cells, because *P. aeruginosa* frequently infects lungs. Whereas MDCK cells are Gb3-negative, H1975 cells express Gb3 (Fig. S2). The glycosphingolipid Gb3 has been previously found to be required for LecA-mediated internalization of *P. aeruginosa* (10).In H1975 cells, LecB triggered also Akt activation, in a dose- and time-dependent manner (Figs. S3A – S3D) and in dependence of PI3K (Figs. S3E and S3F, using the pan-PI3K inhibitors Wortmannin and LY294002, and the Akt inhibitor Triciribine). As a further control, we verified that soluble L-fucose, which prevents LecB from engaging with host cell receptors, is able to inhibit LecB-triggered Akt signaling (Figs. S3G and S3H). This demonstrates that LecB binding to fucosylated receptors is necessary to trigger PI3K/Akt signaling and also validates the purity of our LecB preparation.

To identify apical interaction partners of LecB we apically applied LecB-biotin to polarized MDCK cells, lysed them, and precipitated LecB-receptor complexes with streptavidin beads. Mass spectrometry (MS)-analysis revealed 12 profoundly enriched proteins (Table S1), underscoring the property of LecB to bind to multiple receptors. However, this property also prevented us from singling out a receptor that is responsible for LecB-triggered PI3K signaling, since the list included several proteins for which a capacity to elicit PI3K signaling is known (CEACAM1 (25, 26), Mucin-1 (27), ICAM1 (28), podocalyxin (29, 30)).

Taken together, these findings show that after binding to fucosylated receptors at the plasma membrane of epithelial cells, LecB triggers a Src – PI3K/Akt signaling cascade, which replicates the cellular responses that were observed after binding of live *P. aeruginosa* to apical membranes (13).

### Coating of beads with LecB and expression of LecB by P. aeruginosa enhances their apical uptake

To more realistically model the geometry during infection with *P. aeruginosa*, we utilized bacteria-sized beads that were coated with LecB. In pilot experiments using cell fixation, LecB-coated beads were seen to bind to apical plasma membrane of polarized MDCK cells and to cause local apical accumulation of PH-Akt-GFP/PIP_3_ (Fig. 2A), and many beads were found to be completely internalized by cells (Fig. 2B). Live cell microscopy experiments revealed that apical PH-Akt-GFP/PIP_3_ accumulation is a transient event that occurs before apical uptake of beads by MDCK cells (Fig. 2C, see also Supplementary Movie 1). A detailed quantification showed that biotin-coated control beads were able to trigger apical PH-Akt-GFP/PIP_3_-bursts to some extent, but at a much lower rate than LecB-coated beads (Fig. 2D). Interestingly, PH-Akt-GFP/PIP_3_-bursts caused by control beads were hardly sufficient to mediate cellular uptake, whereas LecB-coated beads were taken up extensively (Fig. 2E). In addition, LecB treatment stimulated macropinocytotic uptake of dextran in H1975 cells (Figs. S4A and S4B), which provides further evidence that LecB activates cellular uptake mechanisms.

**Fig. 2:**
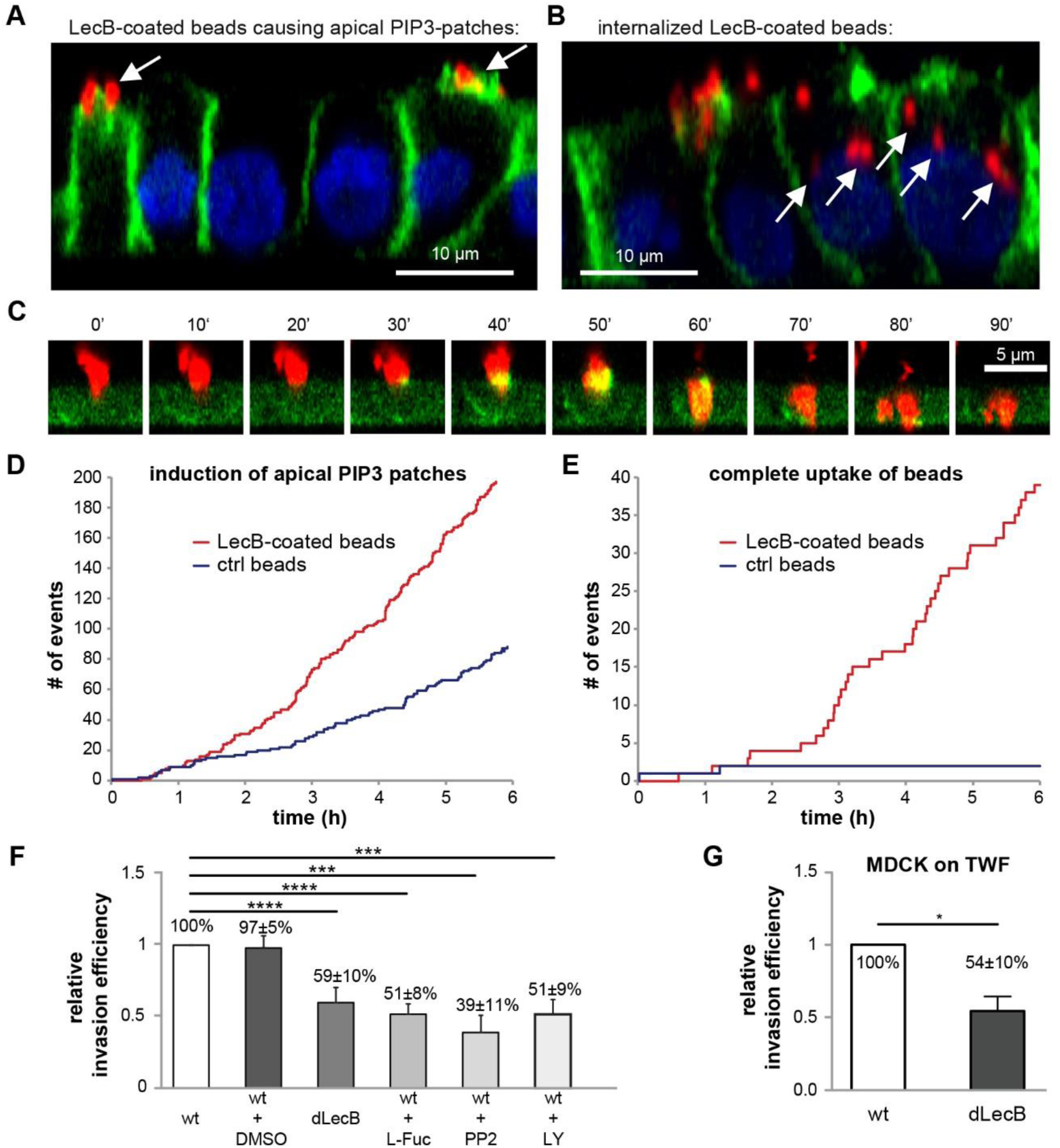
LecB facilitates apical uptake of beads and apical invasion of *P. aeruginosa*. (A) – (B) Red fluorescent LecB-coated bacteria-sized beads with 1 µm diameter were apically applied to MDCK cells stably expressing the PIP_3_ marker PH-Akt-GFP (green) for 6 h; nuclei were stained with DAPI (blue). (A) shows instances of beads causing apical PIP_3_ patches (white arrows), whereas in (B) fully internalized beads are depicted (white arrows). (C) – (E) MDCK cells stably expressing PH-Akt-GFP (green) were allowed to polarize on cover glasses. Red fluorescent beads of 1 µm diameter coated with LecB were applied and live cell confocal imaging was performed. The images show apico-basal cross-sections extracted from confocal image stacks. In (D) the number of induced apical PIP_3_-patches and in (E) the number of beads that are completely taken up over time is depicted for biotin coated beads (ctrl) and LecB-coated beads. (F) Using an amikacin protection assay the invasion efficiencies of wild type (wt) and LecB-deficient (dLecB) PAO1 applied at a MOI of 50 for 2 h on the apical side of polarized MDCK cells grown in 24 well plates were determined. In addition, the invasion efficiencies for bacteria pre-incubated with 100 mg/ml L-fucose (L-Fuc) and for cells treated with PP2 (10 µM) and LY294002 (LY; 10 µM) were measured. Mean values and SEM from n = 8 experiments are shown. (G) Amikacin protection assays measuring the apical invasion of PAO1-wt PAO1-dLecB in MDCK cells grown on transwell filters. Invasion for 2 h, MOI 50, n = 3.

Motivated by these results we investigated if expression of LecB influences host cell uptake of live *P. aeruginosa* bacteria. Indeed, abrogation of LecB expression in *P. aeruginosa* (dLecB) and blockage of LecB with L-fucose diminished the apical uptake of *P. aeruginosa* in polarized MDCK cells (Fig. 2F). In accordance with previous studies (13, 31), inhibition of Src kinases with PP2 and inhibition of PI3K with LY294002 also decreased *P. aeruginosa* uptake (Fig. 2F). Due to the easier handling, the experiments in Fig. 2F were carried out with MDCK cells grown in 24 well plates. For verification, we repeated them with transwell filter-grown MDCK cells, which yielded comparable results (Fig. 2G). Of note, the association of wt and dLecB *P. aeruginosa* with polarized MDCK cells was not significantly different (Fig. S5), which suggests that the observed decrease of invasion efficiency upon deletion of LecB was due to LecB-mediated signaling and not due to reduced host cell binding. In H1975 cells uptake of *P. aeruginosa* was also lowered by LecB deletion (Fig. S4C) and L-fucose treatment (Fig. S4D).

Taken together, these data demonstrate that LecB promotes the uptake of *P. aeruginosa* in polarized epithelial cells from the apical side.

### LecB-mediated PI3K signaling leads to Rac activation and actin rearrangement

To understand the cellular response upon apical LecB stimulation better, we investigated how PI3K activation is linked to *P. aeruginosa* uptake. Motivated by the know correlations between PI3K and Rac activation (32) and the reported implication of Rac in *P. aeruginosa* internalization (31), we carried out experiments using Rac123-G-LISA-assays to test the capability of LecB to activate Rac. We found that apically applied LecB activated Rac in a time-dependent manner in MDCK cells (Fig. 3A) and also in H1975 cells (Fig. 3B). The PI3K inhibitor Wortmannin blocked LecB-mediated Rac activation (Fig. 3C), indicating that PI3K activation occurred upstream of Rac activation.

**Fig. 3:**
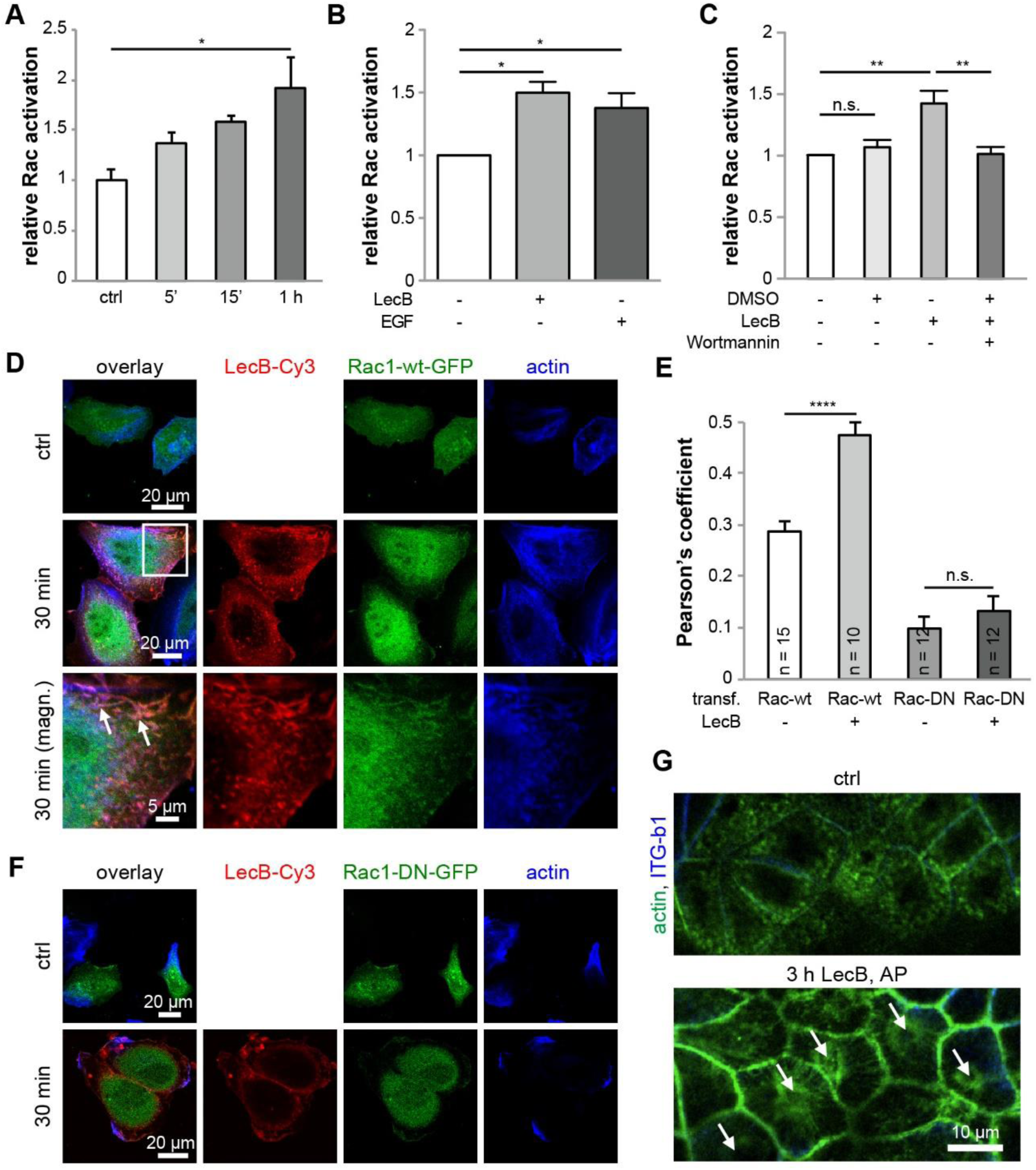
Apical LecB stimulation leads to Rac activation and actin rearrangement. (A) The activation of Rac upon apical LecB treatment of MDCK cells was measured using a Rac123-G-LISA assay; n = 3. (B) H1975 cells were treated with LecB or EGF (20 nM) and Rac activation was measured using a Rac123-G-LISA assay; n = 3. (C) H1975 cells were treated with LecB and Wortmannin (100 nM) and Rac activation was measured using a Rac123-G-LISA assay; n = 6. (D) – (H) H1975 cells transfected with Rac1-wt-GFP (D; green) or Rac1-DN-GFP (F; green) were treated with LecB-Cy3 (red) as indicated, fixed, and stained for actin with phalloidin-Atto647 (blue). In (D) white arrows point to ruffle-like structures where LecB, Rac1-wt-GFP, and actin co-localized. (E) The Pearson’s co-localization coefficient between Rac1-wt-GFP or Rac1-DN-GFP and actin in cells untreated or treated with LecB-Cy3 was determined in individual cells and the average was calculated. (G) MDCK cells treated with LecB as indicated were fixed and stained with phalloidin-Atto488 to stain actin (green) and β1-integrin (blue). Lateral confocal cross-sections along the apical poles of the cells are displayed.

To investigate the consequences of LecB-mediated Rac activation on the actin cytoskeleton further, we utilized unpolarized H1975. The reason for this is that this allowed us to use overexpression of dominant negative (DN) Rac1, which would result in unwanted side effects in polarized MDCK cells, because Rac1 has also roles during polarization of MDCK cells (33). In sparsely seeded H1975 cells LecB caused ruffle-like structures (Fig. 3D), and LecB co-localized with transfected Rac1-wt-GFP and actin in the ruffle-like regions (Fig. 3D, white arrows). To verify that LecB induced recruitment of Rac1-wt-GFP towards actin, we determined the Pearson’s co-localization coefficient between Rac1-wt-GFP and actin, which significantly increased in LecB-treated cells (Fig. 3E). This was not the case when dominant negative (DN) Rac1-GFP (Rac1-DN-GFP) was overexpressed in H1975 cells (Figs. 3F and 3E), showing the requirement of functional Rac for this effect. For verification, we repeated the experiment in untransfected H1975 cells using antibodies recognizing endogenous Rac1 (Fig. S6). Consistently, a recruitment of Rac to actin upon LecB stimulation occurred as well in this experiment.

Apical application of LecB led also to substantial rearrangement of actin at the apical cell pole of MDCK cells (Fig. 3G). In untreated cells, dotted structures representing microvilli and the central actin-devoid region of the periciliary membrane and the primary cilium (34–36) were visible. In cells treated apically for 3 h with LecB, this sub-apical organization of the actin cytoskeleton was completely lost. Actin was recruited to lateral aspects of the cell membrane and actin stress fibers constricting around the central position of the outgrowth of the primary cilium (Fig. 3G, white arrows) appeared.

In summary, these experiments show that LecB-triggered PI3K signaling leads to Rac activation and actin rearrangement. All these processes have been previously observed during internalization of *P. aeruginosa* (22, 31, 37), thus further underscoring the role of LecB for *P. aeruginosa* host cell invasion.

### Apical LecB treatment reversibly removes primary cilia

Motivated by our observation that LecB treatment led to formation of actin stress fibers that appeared to constrict around the basis of the primary cilium, we investigated the effects of LecB on the primary cilium. Interestingly, apical application of LecB removed primary cilia from polarized MDCK cells (Figs. 4A and 4B) within 12 h. This effect was reversible after washout of LecB (Figs. 4C and 4D). Although the potential physiological consequences of primary cilia loss during *P. aeruginosa* infection remain to be investigated, this finding underscores the massive extent of LecB-mediated actin rearrangement.

**Fig. 4:**
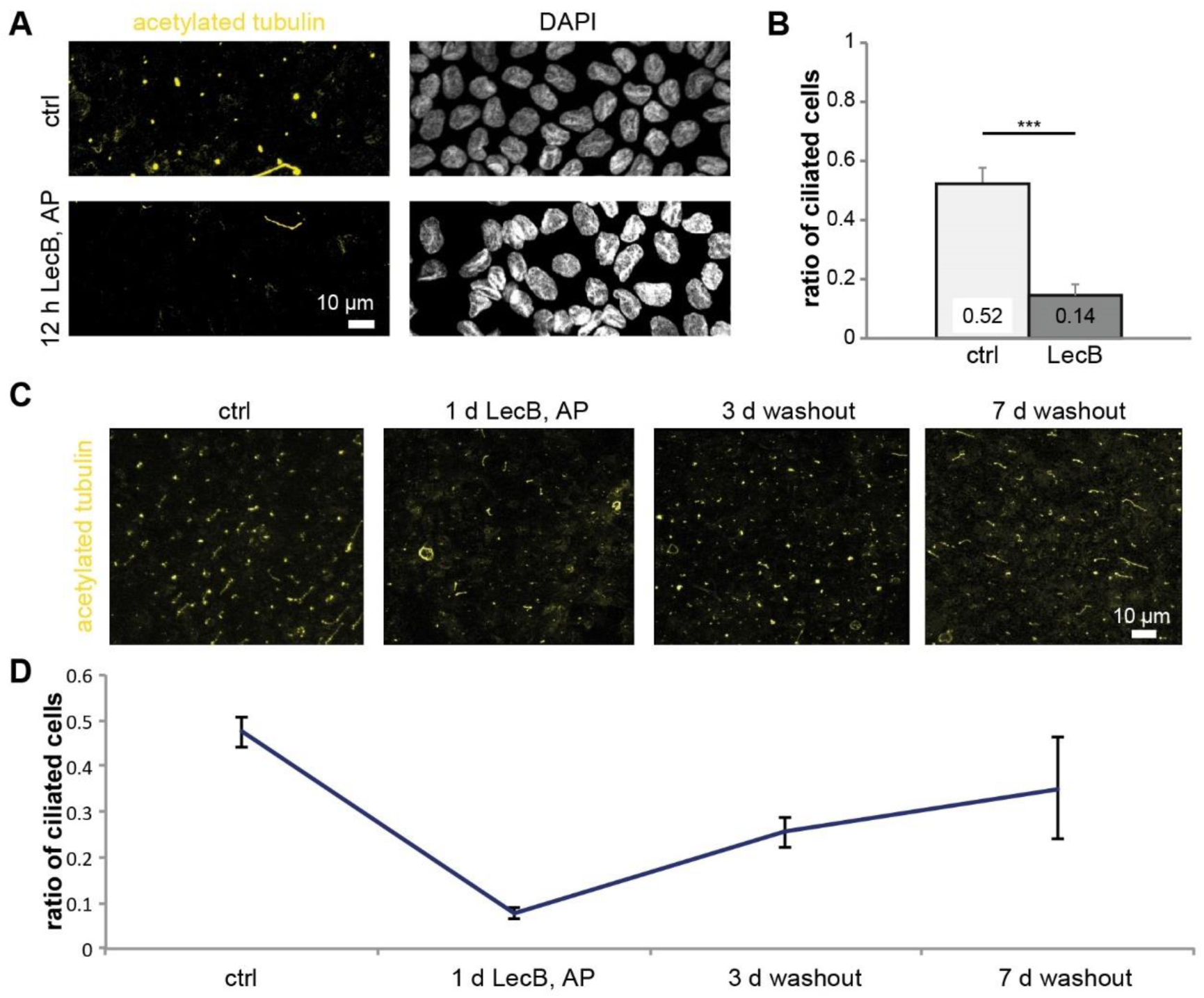
Apical treatment with LecB removes primary cilia in a reversible manner. (A) – (D) MDCK cells were grown on glass coverslips for 10 d. After the indicated treatments, cells were fixed and immunofluorescence staining was performed for acetylated tubulin (yellow) to visualize primary cilia. (A) Nuclei were additionally stained with DAPI (white). Maximum intensity projections of confocal image stacks covering total cell heights are shown. (B) The ratio of ciliated cells was calculated by dividing the number of visible cilia by the total number of cells. 5 fields of view (125 µm x 125 µm) were summed up for n = 1 and the results from n = 3 independent experiments were averaged. (C) MDCK cells were treated with LecB followed by washout as indicated. (D) Quantification of the experiment in (C).

### LecB triggers a feedback loop between caveolin-1 recruitment and PI3K activation

Interestingly, we also found caveolin-1 in the MS screen of LecB interactors (Table S1). Since caveolin-1 is a cytosolic protein, it presumably co-precipitated with LecB-interacting receptors. Motivated by this finding, we further investigated the behavior of caveolin-1 after LecB treatment. In undisturbed MDCK cells caveolin-1 preferentially localized to the basolateral plasma membrane as observed before (38) (Fig. 5A). However, apical LecB treatment resulted in abnormal recruitment of caveolin-1 towards the apical cell pole (Fig. 5A). In addition, the recruitment of caveolin-1 to LecB-receptor complexes was verifiable by WB and increased in a time-dependent manner (Fig. 5B). Interestingly, blocking Src kinases with SU6656 or PP2 and blocking PI3K with LY294002 diminished the co-precipitation of caveolin-1 in complexes with LecB-biotin (Fig. 5C). To directly investigate the requirement of caveolin-1 for LecB-mediated PI3K activation, we knocked down caveolin-1 in MDCK cells using shRNA (Fig. S7). Caveolin-1 knock down almost completely suppressed PI3K activation upon LecB treatment (Fig. 5D).

**Fig. 5:**
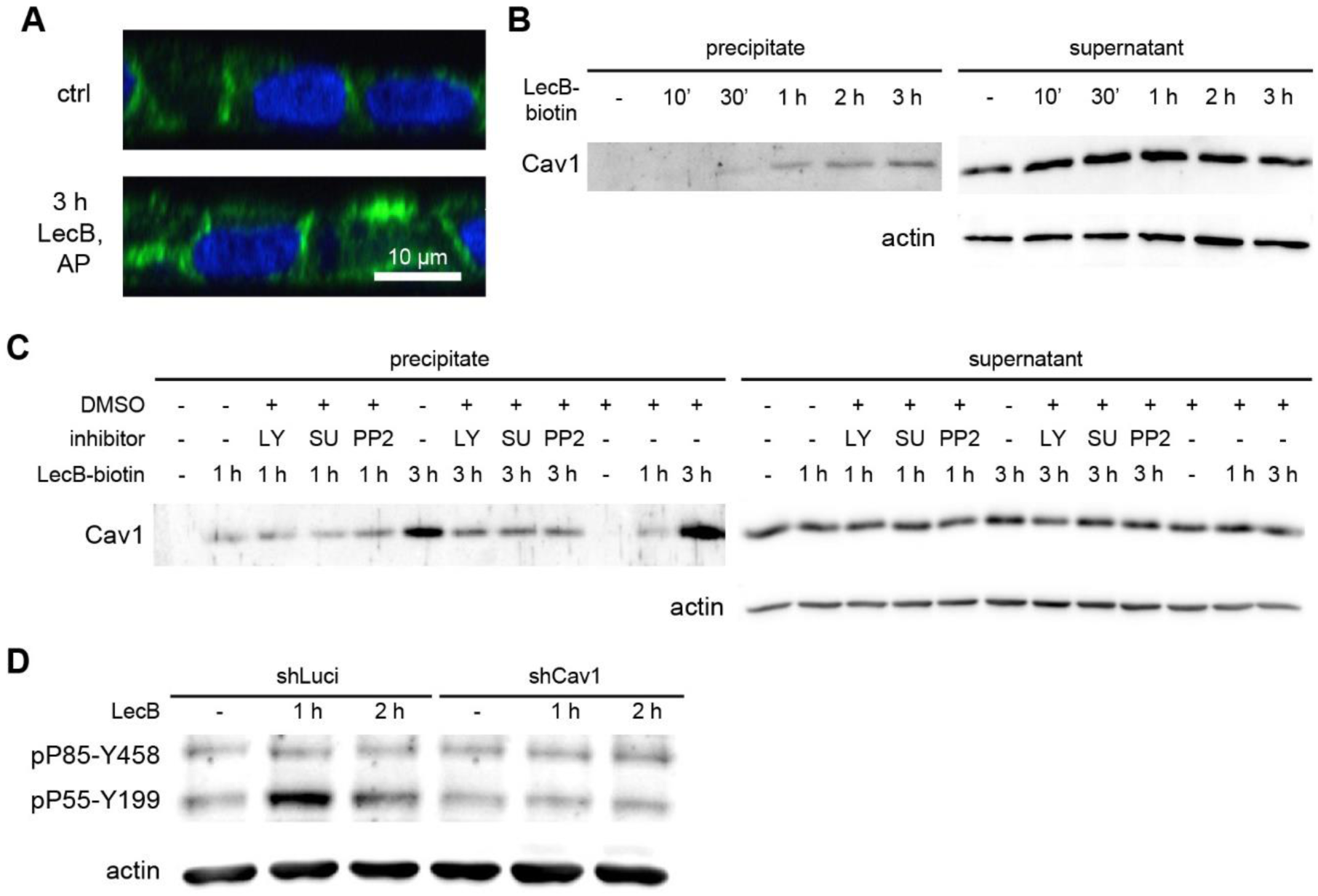
Caveolin-1 is essential for LecB-triggered PI3K signaling. (A) Polarized MDCK cells were treated apically (AP) with LecB as indicated, fixed, and stained for caveolin-1 (green); nuclei were stained with DAPI (blue). (B) – (C) LecB-biotin was apically applied to polarized MDCK cells for the indicated times. After cell lysis, LecB-biotin-receptor complexes were precipitated with streptavidin beads and the precipitate and the supernatant were probed by WB for caveolin-1. (C) Cells were additionally treated with LY294002 (LY; 10 µM), PP2 (10 µM), or SU6656 (SU; 10 µM). (D) Polarized MDCK cells expressing a control shRNA (shLuci) and caveolin-1 knockdown MDCK cells (shCav1) were treated apically with LecB as indicated and subjected to WB analysis using an antibody recognizing active PI3K (pP85-Y458 and pP55-Y199).

Taken together, these data demonstrate that caveolin-1 is apically recruited by LecB stimulation and that this recruitment requires activation of Src kinases and PI3K, whereas caveolin-1 is also required for LecB-triggered PI3K activation. This constitutes a positive feedback loop between caveolin-1 recruitment and PI3K activation.

## Discussion

Here, we demonstrate that LecB is able to trigger a Src-PI3K-Rac signaling cascade, which is modulated by caveolin-1 and leads to actin rearrangement and protrusion formation in order to promote cellular uptake of *P. aeruginosa* bacteria. This adds LecB-triggered signaling to the growing list of *P. aeruginosa* host cell invasion mechanisms, which provokes the question why this bacterium has evolved so many invasion mechanisms and how LecB fits in.

The multitude of invasion mechanisms might be routed in the adaptability of this opportunistic pathogen. *P. aeruginosa* can infect the respiratory tract, urinary tract, eye, and skin (39), and it was demonstrated that this bacterium can invade epithelial cells from the lung (9), cornea (2) and kidneys (22, 40). Considering this diversity, it makes sense that *P. aeruginosa* possesses many invasion mechanisms, which might be used by the bacterium in dependence of the encountered host cell type. One example is lipid zipper-type invasion, which requires the interaction between LecA from *P. aeruginosa* and the glycosphingolipid Gb3 as host cell factor (10). However, this lipid is not expressed in all epithelial cell types. For example the MDCK cells used in this study do not express Gb3 (Fig. S2 and (41)). Nevertheless, *P. aeruginosa* successfully invaded MDCK cells and thus uses alternative pathways, such as LecB-mediated signaling as we demonstrated here. In addition, we show that LecB deletion in *P. aeruginosa* decreased the invasion efficiency also in H1975 cells, which we identified as Gb3-positive (Fig. S2), and it has been demonstrated previously that Gb3 expression in MDCK cells increased the invasion efficiency (10). These examples suggest that invasion mechanisms, such as LecA- and LecB-dependent invasion, are not exclusive but rather function in an additive manner. Our data provides an additional line of evidence for a cooperative function of invasion mechanisms. Coating of bacteria-sized beads with LecB markedly stimulated their uptake into cells, thus demonstrating that LecB alone is sufficient for stimulating cellular uptake. But LecB deletion or LecB blocking with L-fucose did not decrease internalization of *P. aeruginosa* bacteria to the same extent as inhibition of Src kinases and PI3K did. This hints to other bacterial factors that are also able to cause PI3K-dependent uptake into host cells. A potential candidate is type IV pili, since deletion of pili led to a small but significant reduction of PI3K/Akt activation upon apical application of *P. aeruginosa* to polarized Calu-3 cells (12).

How is LecB able to trigger the Src-PI3K-Rac-actin signaling cascade? By MS analysis we showed that LecB binds multiple apical receptors capable of triggering PI3K-signaling: CEACAM1 (25, 26), Mucin-1 (27), ICAM1 (28), and podocalyxin (29, 30). This makes it on one hand more robust for the bacterium to trigger the desired response, but it makes it also difficult for us to isolate a detailed mechanistic picture of LecB action at the apical cell membrane. We hypothesize that LecB has, due to being a tetramer that offers four binding sites (42), the capacity to cross-link and cluster different receptors (19, 43), which is a general mechanism to activate receptor-mediated signaling cascades at the cell membrane. The data we present here provides two independent lines of evidence for this hypothesis. The first line derives from our control experiments with the lectin UEA-I. UEA-I is also able to bind fucose, but it has only two binding sites (44). This makes UEA-I a less ideal cross-linker than the tetrameric LecB, which was shown to be capable of cross-linking fucosylated lipids and integrins (19, 43). Consequently, we found that UEA-I was not capable of eliciting PI3K signaling. This confirms that binding to fucosylated receptors is not enough and additional cross-linking, as in the case of LecB, is required for triggering PI3K signaling. The second line of evidence can be deduced from our experiments regarding caveolin-1. It has been shown that receptor cross-linking is sufficient to aberrantly induce caveolin-1 – containing caveolae at the apical plasma membrane of epithelial cells (38, 45). LecB application at the apical plasma membrane caused also the abnormal recruitment of caveolin-1 to the apical plasma membrane, which can be explained by assuming that LecB cross-linked receptors. In addition, the fact that caveolin-1 knockdown abrogated LecB-mediated PI3K activation together with our finding that caveolin-1 recruitment could be blocked by PI3K inhibitors suggests that there exists a positive feedback loop between PI3K activation and caveolin-1 recruitment. This is strongly supported by our observation that caveolin-1 co-precipitation with apical LecB receptors increased in a time-dependent manner. This also offers an explanation for the previously reported role of caveolin-1 for *P. aeruginosa* host cell invasion (9).

There has been speculation in literature about the initial events that trigger the basolateral patch formation at the apical membrane by *P. aeruginosa* and two possible hypotheses were offered (11): Either membrane damage could be responsible, or a still unknown bacterial factor causes the required PI3K activation. Our results favor the second hypothesis. Binding and cross-linking of apical receptors by LecB offer a direct explanation for PI3K activation and thus identifies LecB as the unknown bacterial factor. In addition, we previously reported that application of purified LecB to the apical plasma membrane of MDCK cells does not induce membrane damage as measured by trypan blue assays that used the fluorescence of trypan blue as sensitive readout (19). Likewise, tight junction integrity was not affected by apical application of LecB (19). This is in agreement with the finding by others that the formation of PIP_3_-rich protrusions during infection with *P. aeruginosa* did not compromise tight junctions (11). This finding also excludes the possibility that LecB-triggered apical PIP_3_ accumulation occurred by diffusive spreading of PIP_3_ from the basolateral plasma membrane, and additionally proves that apical PIP_3_ accumulation was due to LecB-mediated local PI3K activity at the apical plasma membrane.

The involvement of Rac1 for *P. aeruginosa* internalization through the LecB-triggered cascade we describe here will need further clarification. Specifically, the *P. aeruginosa* exotoxin S and exotoxin T are known to contain N-terminal RhoGTPase Activating Protein (RhoGAP) domains, which can hydrolyze GTP to GDP in Rho, Rac, and Cdc42, that leads to cytoskeletal depolymerization and counteracts host cell invasion (46, 47). It will be interesting to investigate if varying expression levels of LecB, exotoxin S, and exotoxin T cause more or less invasive behavior of *P. aeruoginosa*.

In conclusion, our results identify LecB as novel bacterial factor that promotes uptake of *P. aeruginosa* bacteria from the apical side of epithelial cells. Our data suggests that LecB represents a missing link that provides a unifying explanation for many observations that were made during host cell invasion by *P. aeruginosa*. We revealed that LecB is sufficient to trigger the well-known Src-PI3K-Rac signaling cascade (11), which is required for basolateral patch formation at the apical plasma membrane and host cell invasion. LecB-mediated signaling also provides additional rationales for the previously found implication of caveolin-1 in *P. aeruginosa* invasion (9) since we identified here a LecB-triggered positive feedback loop between PI3K activation and caveolin-1 recruitment to the apical plasma membrane.

## Materials and Methods

### Antibodies, plasmids and reagents

Used antibodies are listed in Table S2. The plasmid pPH-Akt-GFP encoding PH-Akt -GFP was a gift from Tamas Balla (Addgene plasmid # 51465). The plasmids encoding wild type Rac1 tagged with GFP (Rac1-wt-GFP) and Rac1-T17N tagged with GFP (Rac1-DN-GFP) were kindly provided by Stefan Linder (University Hospital Hamburg-Eppendorf, Germany).

Recombinant LecB was produced in Escherichia coli BL21(DE3) cells and purified with affinity columns as previously described (19). LecB and fluorophore-conjugated LecB were used at a concentration of 50 µg/ml (4.3 µM) unless stated otherwise. The B-subunit of Shiga toxin 1 (StxB) recombinantly produced in *Escherichia coli* was from Sigma Aldrich. LY294002, Wortmannin, PP2, SU6656, PIK-75, TGX-221, and Triciribine were from Selleckchem. UEA-I was from Vector Labs. Human EGF, L-fucose (6-Deoxy-L-galactose), and FITC-dextran (70 kDa) were from Sigma Aldrich. Phalloidin-Atto488 and phalloidin-Atto647 were from Atto-Tec.

### Mammalian cell culture, creation of stable cell lines

MDCK strain II cells were cultured in Dulbecco’s Modified Eagle’s Medium (DMEM) supplemented with 5% fetal calf serum (FCS) at 37 °C and 5% CO_2_. H1975 cells were maintained in Roswell Park Memorial Institute (RPMI) 1640 medium supplemented with 10% FCS at 37 °C and 5% CO_2_. For generating polarized MDCK monolayers 3×10^5^ MDCK cells were seeded on transwell filters (12 well format, 0.4 µm pore size, polycarbonate membrane, #3401 from Corning) and cultured for 4 d before experiments. For experiments with H1975 cells, 3×10^4^ cells were seeded per 12 mm glass cover slip placed in a 24 well plate and cultured for 1 day. For the creation the MDCK cell line stably expressing PH-Akt-GFP, cells were transfected with the plasmid pPH-Akt-GFP using Lipofectamine 2000 (Thermo Fisher). After allowing the cells to express the proteins overnight, they were trypsinized and plated sparsely in medium containing 1 mg/ml G418. After single colonies had formed, GFP-positive colonies were extracted with cloning rings. At least 6 colonies were extracted for each cell line, grown on transwell filters for 4 d, fixed and stained against the basolateral marker protein β-catenin and the tight junction marker protein ZO-1 to assay their polarized morphology. Based on these results we chose one colony for each cell line for further experiments.

### Caveolin-1 knock down

To achieve knock down of caveolin-1 in MDCK cells a lentivirus-based shRNA system was used based on the plasmids pCMV-ΔR8.91, pMD2G-VSVG, and pLVTH (48). The plasmid pLVTH was modified using Gibson cloning to encode the target sequence for caveolin-1 knock down 5’-GATGTGATTGCAGAACCAG-3’ (49). As control, a shRNA targeted against luciferase, which is not endogenously expressed in MDCK cells, has been used (target sequence: 5′-CGTACGCGGAATACTTCGA-3′). Lentivirus was produced with HEK 293 T cells, purified with sucrose cushion centrifugation (20% sucrose, 4000 g, 14 h), resuspended in MDCK medium and applied to freshly seeded MDCK cells. To ensure a lentivirus transduction efficiency > 80%, GFP fluorescence was checked after 48 h, since pLVTH also encodes GFP. Knock down efficiency was then verified using WB (Fig. S7).

### Immunofluorescence

Cells were washed two times with phosphate-buffered saline without Ca^2+^ and Mg^2+^ (PBS), and then fixed with 4% (w/v) formaldehyde (FA) for 15 min at room temperature. Samples were treated with 50 mM ammonium chloride for 5 min to quench FA and then permeabilized with a SAPO medium (PBS supplemented with 0.2% (w/v) bovine serum albumin and 0.02% (w/v) saponin) for 30 min. Primary antibodies were diluted in SAPO medium and applied on the samples for 60 min at room temperature. After three washes with PBS, secondary dye-labeled antibodies, and, if required, DAPI and dye-labeled phalloidin, were diluted in SAPO medium and applied to the cells for 30 min at room temperature (details for the used antibodies are listed in Table S2). After 5 washes with PBS, cells were mounted for microscopy using glycerol-based medium supplemented with DABCO (MDCK, (50)) or mowiol-based medium (H1975, (51)).

### Microscopy of fixed cells and live cell experiments

For imaging, an A1R confocal microscope (Nikon) equipped with a 60x oil immersion objective (NA

= 1.49) and laser lines at 405 nm, 488 nm, 561 nm, and 641 nm was utilized. Image acquisition and analysis was performed with NIS-Elements 4.10.04 (Nikon).

For live cell experiments, MDCK cells stably expressing PH-Akt-GFP (uptake of LecB-coated beads) were grown as polarized monolayers for 3 d on Lab-Tek II chambered cover glasses (8 well, #1.5 borosilicate glass). The medium was changed to recording medium (Hank’s balanced salt solution (HBSS) supplemented with 1% FCS, 4.5 g/L glucose, and 20 mM HEPES).

### Western Blot analysis

Before WB analysis, cells were starved in media without FCS (16 h for polarized MDCK cells, 2 h for H1975 cells) and stimulation was also carried out in media without FCS. After stimulation cells were washed twice with PBS and lysed in RIPA buffer (20 mM Tris (pH 8), 0.1% (w/v) SDS, 10% (v/v) glycerol, 13.7 mM NaCl, 2mM EDTA, and 0.5% (w/v) sodium deoxycholate in water), supplemented with protease inhibitors (0.8 μM aprotinin, 11 μM leupeptin, 200 μM pefablock) and phosphatase inhibitor (1 mM sodium orthovanadate). Protein concentrations were analyzed using a BCA assay kit (Pierce). Equal amounts of protein per sample were separated by SDS-PAGE and transferred to a nitrocellulose membrane. The membrane was blocked with tris-buffered saline (TBS) supplemented with 0.1% (v/v) Tween 20 and 3% (w/v) BSA for one hour and incubated with primary and HRP-linked secondary antibodies diluted in the blocking solution. Detection was performed by a chemiluminescence reaction using the Fusion-FX7 Advance imaging system (Peqlab Biotechnologie GmbH). If not indicated otherwise, control samples were treated with the same volume of PBS that was used for dissolving LecB in the LecB-treated samples.

### Rac123-G-LISA

Rac activation was measured with a Rac123-G-LISA assay (absorbance based; Cytoskeleton) and performed according to manufacturer’s protocol. Briefly, cells were serum-starved, stimulated as indicated, and then lysed. The lysates were applied to provided 96 well plates and activated Rac was detected at 490 nm using a plate reader (Tecan Safire). If not indicated otherwise, control samples were treated with the same volume of PBS that was used for dissolving LecB in the LecB-treated samples.

### Bacteria culture and invasion assays

For our experiments we used GFP-tagged *P. aeruginosa* PAO1 wild-type (PAO1-wt) and an in-frame deletion LecB mutant (PAO1-dLecB) that were described previously (52). Bacteria were cultured overnight (approximately 16 h) in LB-Miller medium containing 60 µg/ml gentamicin in a shaker (Thriller, Peqlab) at 37 °C and 650 rpm. The bacteria reached an OD measured at 600 nm of approximately 5.

MDCK cells were allowed to polarize on transwell filters or 24-well plates as indicated. H1975 cells were cultured in 24-well plates to a confluence of 70-80%. Overnight cultures of PAO1-wt and PAO1-dLecB were pelleted and resuspended in DMEM (MDCK) or RPMI (H1975) and incubated for 30 min at 37 °C. For inhibition with L-fucose, 100 mg/ml L-fucose was added during this incubation. The inhibitors PP2 and LY294002 were pre-incubated for 30 min with the cells and kept on the cells during the whole experiment. Next, the concentration of bacteria was adjusted to yield the desired multiplicity of infection (MOI) of 50. For determining the total number of bacteria, cells were incubated with bacteria for 2 h at 37 °C, washed three times with PBS and then lysed with 0.25% (v/v) Triton X-100. Serial dilutions of the cell extracts were made and plated on LB–Miller agar plates containing gentamicin (60 μg/ml) and incubated over night at 37 °C. The number of bacterial colonies was counted on the next day. For determining the number of invading bacteria, cells were incubated with bacteria for 2 h at 37 °C and washed three times with PBS. Then extracellular bacteria were killed by treatment with 400 μg/ml amikacin sulfate (Sigma Aldrich) for 2 h at 37 °C. After lysis with 0.25% (v/v) Triton X-100, bacteria numbers were counted as described before. The invasion efficiencies were calculated by dividing the number of invading bacteria by the total number of bacteria. To enable comparison between different experiments, the invasion efficiencies in a single experiment were normalized to the invasion efficiency of the untreated sample and then the mean value from repeated experiments was calculated.

### Labeling of lectins

LecB was labeled with Cy3 mono-reactive NHS ester (GE Healthcare) or with biotin using NHS-PEG12-biotin (Thermo Fisher) according to the instructions of the manufacturers and purified using PD-10 desalting columns (GE Healthcare). StxB was labeled with NHS-ester conjugated Alexa488 (Thermo Fisher).

### Preparation of LecB-coated beads

Biotinylated LecB (LecB-biotin) was incubated with a solution containing streptavidin-coated polystyrene beads containing the dye Flash red with 1 µm diameter (Bangs Laboratories). To ensure homogenous coverage with LecB-biotin, a ten-fold molar excess of LecB-biotin as compared to the available streptavidin binding sites on the beads was used and then beads were washed three times with PBS. In control beads the streptavidin binding sites were saturated with biotin.

### Mass spectrometry-based identification of LecB interaction partners

MDCK cells were cultured in SILAC media for 9 passages and then seeded on transwell filters and allowed to polarize for 4 d. For the first sample, biotinylated LecB was applied to the apical side of light-SILAC-labeled cells and on the basolateral side of medium-SILAC-labeled cells, whereas heavy-SILAC-labeled cells received no stimulation and served as control. For the second sample, the treatment conditions were permuted. After lysis with IP lysis buffer, the different SILAC lysates were combined and LecB-biotin-receptor complexes were precipitated using streptavidin agarose beads as described before. Eluted LecB-biotin-receptor complexes were then prepared for MS analysis using SDS-PAGE gel electrophoresis. Gels were cut into pieces, proteins therein digested with trypsin and resulting peptides were purified by STAGE tips. MS analysis was carried out as described previously (19) using a 1200 HPLC (Agilent Technologies, Waldbronn, Germany) connected online to a LTQ Orbitrap XL mass spectrometer (Thermo Fisher Scientific, Bremen, Germany). From the generated list of MS-identified proteins we defined those proteins as LecB interaction partners that showed more than 2-fold enrichment on a log2-scale over controls in both SILAC samples (see Table S1).

### Statistics

If not stated otherwise, data obtained from n = 3 independent experiments were used the calculate arithmetic means and error bars represent standard error mean (SEM). Statistical significance analysis was carried out using GraphPad Prism 5. For determining the significance in experiments with multiple conditions one-way ANOVA with Bonferroni’s post hoc testing was applied. For determining the significance in experiments in which values were measured for one condition relative to the control condition, one sample t-testing was applied. n.s. denotes not significant, * denotes P < 0.05, ** denotes P < 0.01, *** denotes P < 0.001, and **** denotes P < 0.0001. All primary data are available from the authors upon request.

## Acknowledgements

This work was supported by the German Research Foundation grants [RO 4341/2-1] and [Major Research Instrumentation (project number: 438033605)], the Excellence Initiative of the German Research Foundation [EXC 294 and EXC 2189], the Ministry of Science, Research and the Arts of Baden-Württemberg [Az: 33-7532.20], the Freiburg Institute for Advanced Studies (FRIAS) and a starting grant from the European Research Council [Programme “Ideas,” ERC-2011-StG 282105]. R.T. acknowledges support from the Ministry of Science, Research and the Arts of Baden-Württemberg [Az: 7533-30-10/25/36]. E.W.K. acknowledges support from the German Research Foundation [KFO 201, KU 1504/5-1, and SFB1140].

## Supplementary Data Legends

**Fig. S1: Control experiments related to Figure 1**

(A) MDCK cells were treated apically with LecB and with the indicated concentrations of the p110α-specific inhibitor PIK-75 for 1 h and Akt activation (pAkt-S473) was probed by WB analysis. (B) MDCK cells were treated apically with LecB and with the indicated concentrations of the p110β-specific inhibitor TGX-221 for 1 h and Akt activation (pAkt-S473) was probed by WB analysis. (C) MDCK cells were treated apically with 50 µg/ml UEA-I or LecB and Akt activation (pAkt-S473) was probed by WB analysis.

**Fig. S2: Evaluation of Gb3 expression in MDCK cells and H1975 cells**

MDCK cells (A) and H1975 cells (B) were seeded sparsely on glass cover slips and then incubated for 30 min at 37°C with 1 µg/ml StxB-Alexa488 (green). StxB is a lectin that specifically binds the glycosphingolipid Gb3. After fixation, cell nuclei were stained with DAPI (blue) and all samples were imaged with a confocal microscope using the same settings to ensure comparability. For each cell type, three different randomly chosen regions of interest (ROI) are displayed. Whereas MDCK cells do not bind StxB and are therefore Gb3-negative, H1975 cells show detectable binding of StxB and are hence expressing Gb3.

**Fig. S3: In H1975 lung epithelial cells LecB also activates PI3K/Akt signaling**

(A) – (C) H1975 cells were treated with LecB as indicated and Akt activation (pAkt-S473) was probed by WB analysis. The image depicts a representative WB; quantifications from n = 3 independent experiments for the dose-dependence and the time-dependence are depicted in (B) and (C), respectively. (D) H1975 cells were treated with LecB-Cy3 (red), fixed, and activated Akt was visualized by an antibody specific for pAkt-S473 (green). (E) – (F) H1975 cells were treated with LecB and PI3K-inhibitors Wortmannin (100 nM), LY294002 (10 µM), and the Akt inhibitor Triciribine (10 µM) for 1 h and Akt activation (pAkt-S473) was probed by WB analysis. (E) shows a representative WB, a quantification from n = 3 independent experiments is depicted in (F). (G) – (H) H1975 cells were treated with LecB and L-fucose (43 mM) for 1 h and Akt activation (pAkt-S473) was probed by WB analysis. (G) shows a representative WB, a quantification from n = 3 independent experiments is depicted in (H).

**Fig. S4: LecB stimulates macropinocytosis and facilitates *P. aeruginosa* invasion in H1975 cells**

(A) – (B) Representative images of untreated H1975 cells and cells treated with LecB-Cy3 for 2 h (red). To probe macropinocytosis cells were also incubated with 0.2 µM FITC-dextran (green); nuclei were stained with DAPI (blue). White arrows point to internalized FITC-dextran co-localizing with LecB-Cy3. (D) Quantification of FITC-dextran uptake in untreated and LecB-treated cells; n = 3. (C) – (D) Amikacin protection assays measuring the invasion of PAO1-wt and PAO1-dLecB (C) and PAO1-wt pre-incubated with 100 mg/ml L-fucose (D) in H1975 cells. Invasion for 2 h, MOI 50, n = 3.

**Fig. S5: Comparison of cell association of wt and dLecB *P*.*aeruginosa***

Wt and dLecB *P. aeruginosa* were applied to polarized MDCK cells at MOI 50 for 2 h. Afterwards, cells were lysed with 0.25% (v/v) Triton X-100. Serial dilutions of the cell extracts were made and plated on LB–Miller agar plates containing gentamicin (60 μg/ml) for counting. This corresponds to the procedure for determining the total number of bacteria in the amikacin protection assays. The graph shows the mean values of counted bacteria from n = 8 experiments.

**Fig. S6: Control experiment related to Figure 3**

(A) – (B) H1975 cells were treated with LecB-Cy3 (red) as indicated, fixed, and stained for actin with phalloidin-Atto647 (blue) and endogenous Rac1 (green) and then imaged with a confocal microscope. (A) Representative images. (B) The Pearson’s co-localization coefficient between endogenous Rac1 and actin was determined in individual cells and the average was calculated.

**Fig. S7: Verification of caveolin-1 knockdown in shCav1 cells**

(A) – (B) The expression of caveolin-1 in caveolin-1 knockdown MDCK cells (shCav1) and wild type MDCK cells (wt) were assessed by WB analysis. (A) Representative WB. (B) Quantification of caveolin-1 knockdown from n = 3 independent experiments. To calculate the normalized caveolin-1 expression, the caveolin-1 band intensities were divided by actin band intensities for each individual sample and averaged. Error bars represent SEM, statistical significance was evaluated by a paired two-sided t-test, ** denotes p <0.01.

**Table S1: List of apical LecB interaction partners identified by SILAC MS**

**Table S2: Lists of used primary and secondary antibodies** (WB … Western Blot, IF … immunofluorescence)

**Supplementary Movie 1:**

The movie shows a time lapse of the internalization of a LecB-coated bead cluster (red) into polarized MDCK cells expressing PH-Akt-GFP (green). The time between individual frames is 10 min and the movie is replayed at 3 frames per second.

